# Validation of a library of cGMP-compliant human pluripotent stem cell lines for use in liver therapy

**DOI:** 10.1101/298760

**Authors:** Samuel J I Blackford, Soon Seng Ng, Joe M Segal, Aileen J F King, Jennifer Moore, Michael Sheldon, Dusko Ilic, Anil Dhawan, Ragai Mitry, S Tamir Rashid

**Affiliations:** Centre for Stem Cells and Regenerative Medicine, King’s College London, England, UK; Institute for Liver Studies, King’s College Hospital, King’s College London, England, UK; Diabetes Research Group, Faculty of Life Sciences & Medicine, King’s College London, England, UK; RUCDR Infinite Biologics, Rutgers University, New Brunswick, NJ, USA; Stem Cell Laboratory, Department of Women and Children’s Health, Faculty of Life Sciences & Medicine, King’s College London, London, UK

**Keywords:** Pluripotent Stem Cells, Hepatocytes, Bioengineering, Biocompatible Materials

## Abstract

Recent advancements in the production of hepatocytes from human pluripotent stem cells (hPSC-Heps) afford tremendous possibilities for treatment of patients with liver disease. Validated current good manufacturing practice (cGMP) lines are an essential prerequisite for such applications but have only recently been established. Whether such cGMP lines are capable of hepatic differentiation is not known. To address this knowledge gap, we examined the proficiency of three recently derived cGMP lines (two hiPSC and one hESC) to differentiate into hepatocytes and their suitability for therapy. hPSC-Heps generated using a chemically defined four-step hepatic differentiation protocol uniformly demonstrated highly reproducible phenotypes and functionality. Seeding into a 3D PEG-DA fabricated inverted colloid crystal (ICC) scaffold converted these immature progenitors into more advanced hepatic tissue structures. Hepatic constructs could also be successfully encapsulated into the immune-privileged material alginate. This is the first report we are aware of demonstrating cGMP-compliant hPSCs can generate cells with advanced hepatic function potentially suitable for future therapeutic applications.

## Introduction

For patient’s presenting with end-stage chronic liver disease (ESLD) or acute liver failure (ALF), orthotopic transplantation of a donor liver remains the only curative treatment. Due to lengthening waiting lists and a severe scarcity of donors, mortality rates are increasing annually [1]. As a result, an alternative strategy for treating these patients is urgently required. Allogenic transplantation of primary adult hepatocytes, the major functional cell-type of the liver, is considered a viable solution in certain clinical indications for example [2–4]. Lack of sufficient numbers of high-quality hepatocytes; a result of isolating cells from tissue deemed unsuitable for transplantation, has limited the success of this programme [5,6].

Derivation of human embryonic stem cells (hESCs) [7] and their related induced pluripotent stem cells (hiPSCs) [8,9] however has generated growing optimism that the development of cellular therapies, such as would be suitable for liver disease, was finally an obtainable goal [10]. Unlike primary hepatocytes, which cannot be cultured or expanded *in vitro* [11,12], human pluripotent stem cells (hPSCs) possess an unlimited ability for self-renewal [13]. This capability to produce large batches of cells is of clinical significance as hepatocytes numbers approaching 8 billion may be required for transplant when correcting metabolic liver function in pediatric patients [14], or 15 billion to support liver failure in adults [15]. Studies have shown that both hESCs [16,17] and hiPSCs [18–20] can be differentiated into hepatocytes (hPSC-Heps), sharing functional attributes of their *in vivo* equivalents, including albumin / alpha-1 antitrypsin (A1AT) protein secretion, Cytochrome P450 activity and glycogen storage. As research tools, hPSCs have delivered novel insights into human hepatic development [21], the creation of liver disease models [22,23], and provided new platforms for pharmacological testing [24]. Furthermore, the successful, albeit limited, repopulation of rodent livers following transplantation of both hESC, and hiPSC-derived hepatocytes has been reported by several groups [25–27], suggesting hPSC-Heps may be a viable treatment option for patients with liver disease.

The stem cell field has developed at an exceptional rate, which has resulted in the first human trials using cells derived from hESCs/hiPSCs being undertaken [28–32]. While approved for clinical use, these lines were in fact developed for research purposes and not produced under current good manufacturing practice (cGMP) guidelines. These manufacturing regulations for stem cell therapy products are described by the Food and Drug Administration in the United States of America (USA), and since the 2004 European Union Tissues and Cells Directive (EUTCD), the European Medicines Agency within the European Union.

Generating cells under cGMP conditions ensures their clinical safety, and for cellular therapies should be differentiated in fully defined, xeno-free conditions [33] to ensure reproducibility, and prevent xeno-mediated infection or immune rejection [34].

In 2011 scientists from King’s College London submitted the first xeno-free clinical grade hESCs [35] to the UK Stem Cell Bank [36]. More recently, cGMP-compliant hiPSCs have been generated by teams in China [37], Japan [38], USA [39] and the United Kingdom [40]. These clinically compliant lines have been extensively characterized as pluripotent with profiles comparable to previously validated hPSCs derived outside of these manufacturing guidelines [41]. However, given previous reports describing the varied potential for hPSC lines to differentiate into target cell types [42–44], evaluation of their hepatic differentiation potential represents an essential pre-requisite to further clinical development work. The objective of this study was therefore to undertake this evaluation and then test the potential of differentiated cells to be engineered into three-dimensional (3D) constructs suitable for clinical application.

## Materials & Methods

### Cell lines and cell culture

Two cGMP hiPSC lines (CGT-RCiB-10 [Line 1; Cell & Gene Therapy Catapult, London, UK] and LiPSC-GR1.1 [Line 2; Lonza, Walkersville, USA]) and one cGMP hESC line (KCL037 [Line 3; Gifted from D. Ilic, King’s College London]) were used in this study. All three lines were recovered into the culture conditions recommended by their respective suppliers. After two passages, each of the lines was subsequently maintained on Vitronectin XF (STEMCELL Technologies, Vancouver, Canada) coated tissue culture plastic in TeSR-E8 (STEMCELL Technologies, Vancouver, Canada) and passaged every 4 days using Gentle Cell Dissociation Reagent (STEMCELL Technologies, Vancouver, Canada). Line 3 was passaged in TeSR-8 supplemented with 10μM Y-27632 dihydrochloride (R&D Systems, Minneapolis, USA) to ensure cell survival.

Hepatocyte differentiation was carried out in Essential 6 Medium (Thermo Fisher Scientific, Waltham, USA) [Days 1-2], RPMI-1640 Medium (Sigma-Aldrich, St. Louis, USA) [Days3-8] and HepatoZYME-SFM (Thermo Fisher Scientific, Waltham, USA) [Day 9 onwards]. Day 21 hPSC-Heps were dissociated into a single-cell suspension using TrypLE Express Enzyme (1X), no phenol red (Thermo Fisher Scientific, Waltham, USA).

### Brightfield and immunofluorescence imaging

Brightfield microscopy was performed on a Leica DMIL LED inverted microscope and imaged using the Leica DFC3000 G camera (Leica Microsystems, Wetzlar, Germany).

For immunofluorescence staining, samples were fixed for 10 minutes with 4% w/v PFA, and then blocked and permeabilized in 1% w/v bovine serum albumin (Sigma-Aldrich, St. Louis, USA), 3% donkey serum (Thermo Fisher Scientific, Waltham, USA) and 0.1% Triton X-100 (Sigma-Aldrich, St. Louis, USA). An additional 10 minutes of permeabilization was performed using 0.5% Triton X-100 for detection of nuclear antigens. Primary antibodies were applied for 1 hour and after wash steps Alexa Fluor-555/488 conjugated secondary antibodies (Thermo Fisher Scientific, Waltham, USA) were incubated for 40 minutes.

NucBlue Fixed Cell ReadyProdes Reagent (Thermo Fisher Scientific, Waltham, USA) was applied for visualization of cell nuclei. Imaging for two-dimensional (2D) culture was performed on an Operetta High Content Screening System (PerkinElmer, Waltham, USA), and for 3D culture, a Nikon Eclipse Ti inverted spinning disk confocal microscope (Nikon, Minto, Japan).

### Real-Time PCR

Total RNA was isolated using the RNeasy Mini Kit (QIAGEN, Hilden, Germany) according to manufacturer’s protocol. RNA was quantified spectrophotometrically using the NanoDrop 2000 (Thermo Fisher Scientific, Waltham, USA). 350ng of total RNA was used to produce first-strand cDNA using the SuperScript VILO cDNA synthesis kit (Thermo Fisher Scientific, Waltham, USA). Quantitative real-time PCR (RT-PCR) was performed in a 10μl reaction mixture consisting of cDNA, custom designed oligonucleotide primers (Sigma-Aldrich, St. Louis, USA) and Fast SYBR Green PCR Master Mix (Thermo Fisher Scientific, Waltham, USA), on a CFX384 Touch Real-Time PCR Detection System (Bio-Rad, Hercules, USA). ACTB mRNA was used for housekeeping normalization.

### Flow cytometric analysis

Adherent cells were dissociated into a single-cell suspension using TrypLE Express Enzyme (1X), no phenol red, and subsequently treated for 30 minutes with LIVE/DEAD Fixable Cell Stain (Thermo Fisher Scientific, Waltham, USA) and then fixed using 4% w/v PFA for 10 minutes. Cells were incubated with fluorophore-conjugated antibodies for 30 minutes in the dark, and then washed twice with PBS. Immunophenotyping was carried out using the BD FACSCanto II system (Becton Dickinson, Franklin Lakes, USA) and analyzed using FlowJo software (Becton Dickinson, Franklin Lakes, USA).

### Assessment of hepatic function

Albumin production of hPSC-Heps was measured using the Human Albumin Quantification Set (Bethyl Laboratories, Montgomery, USA). Culture medium supernatants were collected after 48 hours and stored at −20°C. Enzyme-linked immunosorbent assay (ELISA) was carried out according to manufacturer’s instructions. Absorbance was measured at 450nm on a Promega GloMax Multi+ Detection System plate reader (Promega, Madison, USA).

Native cytochrome P450 CYP3A4 activity was assessed using the CYP3A4 P450-Glo Assay with Luciferin-IPA (Promega, Madison, USA). The bioluminescent substrate was incubated on hPSC-Heps for 1 hour before being collected for analysis. Luminescence was measured using a Promega GloMax Discover multimode microplate reader (Promega, Madison, USA).

### Fabrication of inverse colloidal crystal (ICC) poly(ethylene glycol)-diacrylate (PEG-DA) scaffolds

Thermo Scientific 4000 Series monosized polystyrene beads of 100±1.5μm diameter (Thermo Fisher Scientific, Waltham, USA) were suspended in 70% EtOH and agitated using an ultrasonic bath. The dispersed bead suspension was seeded into hexagonal polypropylene moulds and left to dry overnight on an orbital shaker.

A self-standing colloidal crystal lattice was produced through annealing the beads at 120°C for 4 hours. PEGDA (Thermo Fisher Scientific, Waltham, USA) acrylate-PEGN-hydroxysuccinimide (Laysan Bio Inc, Arab, USA) and Irgacure 2959 photoinitiator (BASF, ^London, UK) were mixed together in dH_2_0 at a concentration of 50, 10 and 1% w/v^ respectively. The bead lattices were placed within this precursor solution, and centrifugation (500G, 5 minutes) was performed to ensure complete infiltration. Hydrogel fabrication was completed through UV light induced gelation of the precursor solution and the polystyrene crystal lattice was removed from the scaffold through tetrahydrofuran (Sigma-Aldrich, St. Louis, USA) soaking for 4 hours.

### Generation of hPSC-derived hepatocyte spheroids

A single cell suspension of day 21 hPSC-Heps was prepared and 0.3×10^6^ cells were seeded per well of a 24-well Aggrewell-400 (STEMCELL Technologies, Vancouver, Canada). Aggrewell plates were prepared as recommended by the supplier. Centrifugation at 200G for 3 minutes was carried out to deposit cells into the microwells of the plate.

### Alginate encapsulation of hPSC-derived hepatocyte spheroids

Encapsulation was performed as previously published [45,46]. In brief, spheroids were washed in saline before being re-suspended into a final 1.8% ultra-pure low-viscosity, high-glucuronic acid (≥60%), sodium alginate (FMC BioPolymer, Drammen, Norway) solution, which was then delivered by syringe pump through a 0.2mm diameter nozzle, from which droplets were electrostatically deposited into a divalent cationic solution (1mM BaCl2 + 50mM CaCl2) to cause gelation.

### Live/Dead staining

Fluorescine diacete (FDA) (Sigma-Aldrich, St. Louis, USA) and cell-impermeant ethidium homodimer-1 (EthD-1) (Thermo Fisher Scientific, Waltham, USA) were used as recommended by the supplier for staining of viable and dead cells. Spheroids and alginate encapsulated cells were incubated in 4μM EthD-1 for 35 minutes, washed with Hank’s Balanced Salt Solution (HBSS) containing calcium (Thermo Fisher Scientific, Waltham, USA), then incubated in 50μg/ml FDA for 90 seconds, and finally washed 5 times with HBSS before imaging on a Leica TCS SP8 Confocal laser scanning microscope (Leica Microsystems, Wetzlar, Germany).

## Results

We firstly recovered two lines of hiPSCs, as well as one line of hESCs, each of which having been derived independently using cGMP-compliant protocols. We maintained all lines in identical culture conditions comprising of xeno-free cell culture matrix, Vitronectin, and chemically defined pluripotency culture medium, TeSR-E8. After several passages within these culture conditions, each of the lines had fully reconditioned, with comparable cell morphologies and colony sizes (Supplementary data 1); each line producing characteristic rounded colonies, with small densely packed cells. We then began investigating the capability of each line in producing high quality hPSC-Heps using an adapted four-stage hepatocyte differentiation protocol based on Hannan et al. [47], and their potential application in downstream clinical applications by achieving advanced phenotypes in macroporous hydrogels and ensuring viability within transplantation ready cell encapsulation models for ALF therapy (Figure 1).

**Figure 1.**
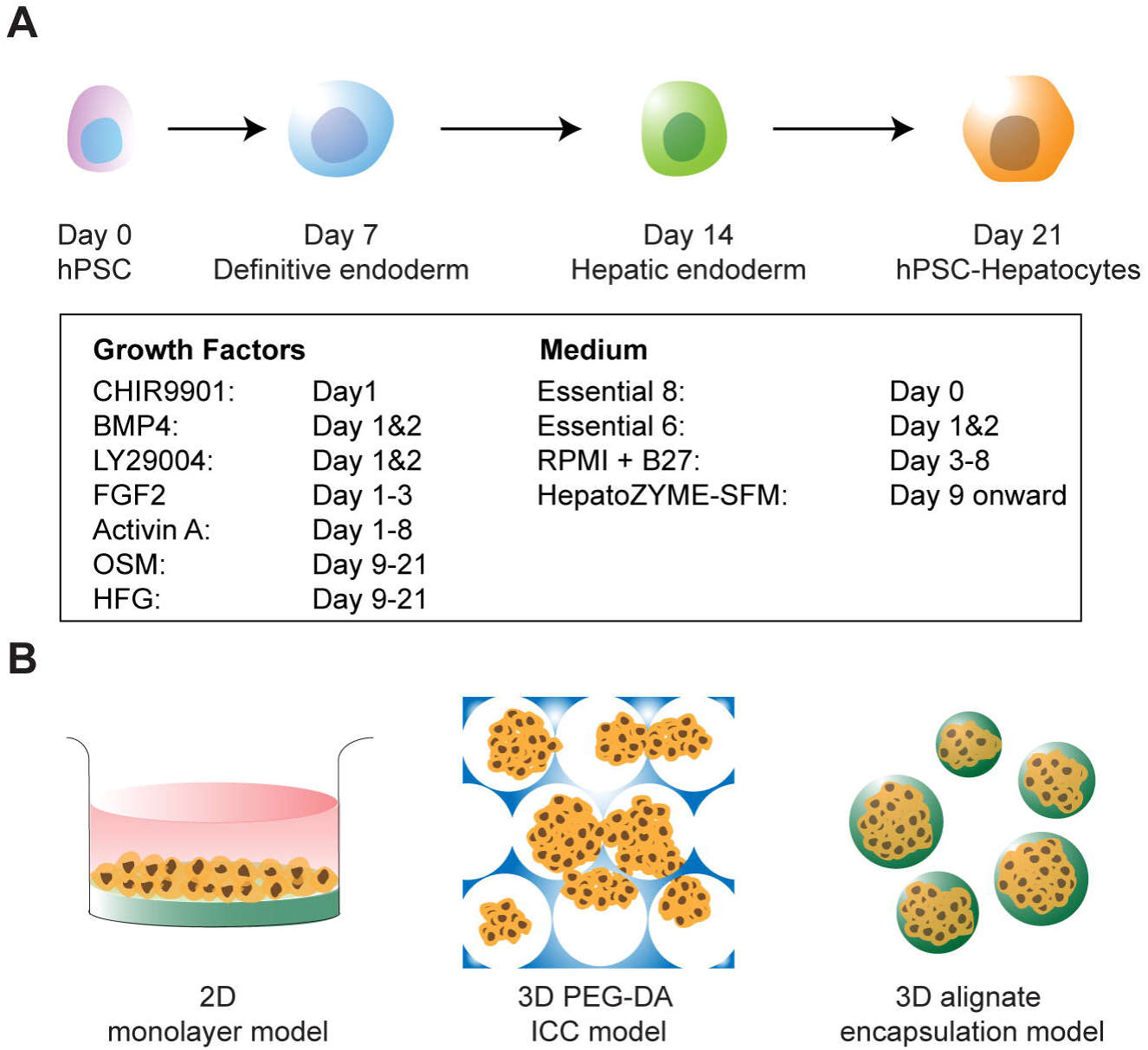
Generation of human pluripotent stem cell-derived hepatocytes and their therapeutic potential in various model systems. (**A**) Four-stage hepatic differentiation from human pluripotent stem cells (hPSC) to definitive endoderm (DE), hepatic endoderm (HE) and subsequently hepatocytes (hPSC-Heps) over a 21-day protocol by a defined cocktail of growth factors and small molecules (in the box). (**B**) Schematic of further maturation of hPSC-Heps in two-dimensional (2D) culture dish coated with Type I collagen, three-dimensional (3D) inverse colloidal crystal scaffold coated with Type I collagen (ICC) and encapsulation within 3D alginate microspheres.

### Differentiation of cGMP-compliant stem cells towards hPSC-derived hepatocytes

To prepare for hepatic differentiation, we seeded fragmented hPSC colonies obtained through enzyme-free dissociation with minimal trituration onto gelatin coated tissue culture dishes. We treated hESCs with the small molecule Y-27632 dihydrochloride, an inhibitor of the RHO/ROCK pathway, overnight to ensure sufficient cell attachment onto the gelatin. We found that the hiPSC lines did not require the addition of this small molecule to be passaged, or adhere to gelatin. We then initiated hepatic differentiation two days post-seeding; having allowed enough time for the hPSCs to reestablish rounded colonies, with the outer cells tending to have a larger, more spread morphology.

We closely monitored the morphology of the cells throughout the differentiation and cross-compared against that of our previous publication, that used non-cGMP-compliant hiPSCs, to ensure appropriate differentiation was achieved (Figure 2A). Upon differentiation, the clear borders signifying stem cell colonies dissipate gradually with the peripheral cells start to spread and migrate out sporadically by day 1 post-differentiation. These protruding cells expand in size and proliferate to close the space between neighboring colonies until a confluent monolayer is formed by day 4. At this stage, cells exhibit definitive endoderm-like morphology which persists until the media condition is changed to contain oncostatin-M and hepatocyte growth factor by day 9. After this alteration, the morphology becomes more dynamic as the cells continue to differentiate. By day 14 the cells start to transform from their elongated morphology, observed at day 11, into a more cuboidal shape. The signature, well defined, polyhedral morphology of hepatocytes is observed across the whole culture by day 17. As demonstrated by the dynamic morphological transformation throughout the course of differentiation, all three of cGMP-compliant lines appear capable of generating populations of definitive endoderm, hepatic endoderm, and subsequently hPSC-Heps (Figure 2B).

**Figure 2.**
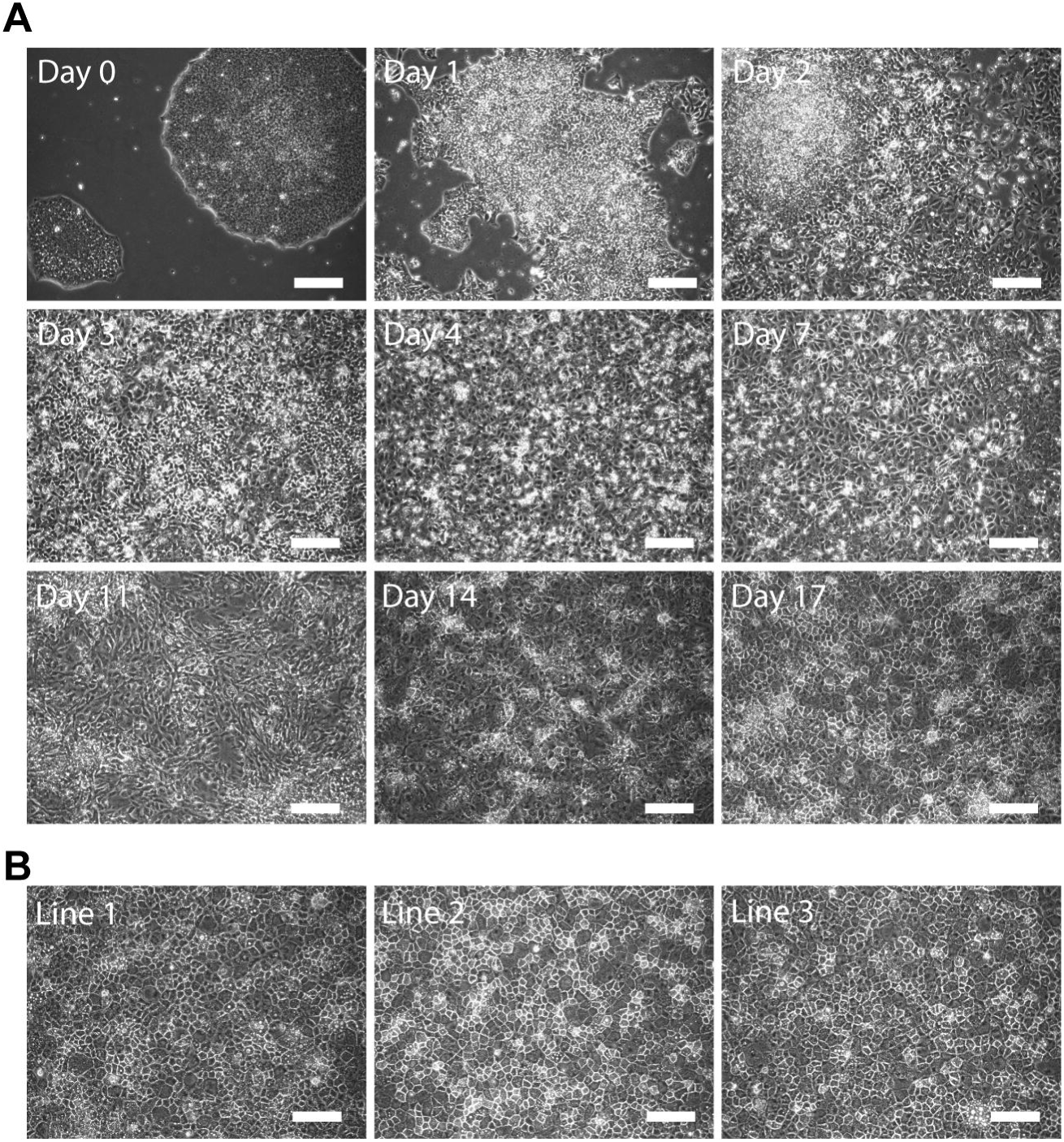
Morphological characterization of hepatic differentiated cells. **(A)** Brightfield microscopy images revealing the morphological transformation from Day 0 pluripotent stem cell colony to Day 17 polyhedral hepatocytes. **(B)** Representative images of day 21 hPSC-Heps generated from three different cGMP-compliant lines. Scale bars: 100μm.

### Characterization of cGMP-compliant hPSC-derived hepatocytes

We next validated that the cGMP-hPSCs were differentiating through the correct developmental lineage trajectory by collecting the cells and quantifying their mRNA expression at four distinct stages of the protocol (Figure 3A). These time-points represent undifferentiated hPSCs, definitive endoderm (DE, day 7), hepatic endoderm (HE, day 14) and finally hPSC-Heps (day 21).

**Figure 3.**
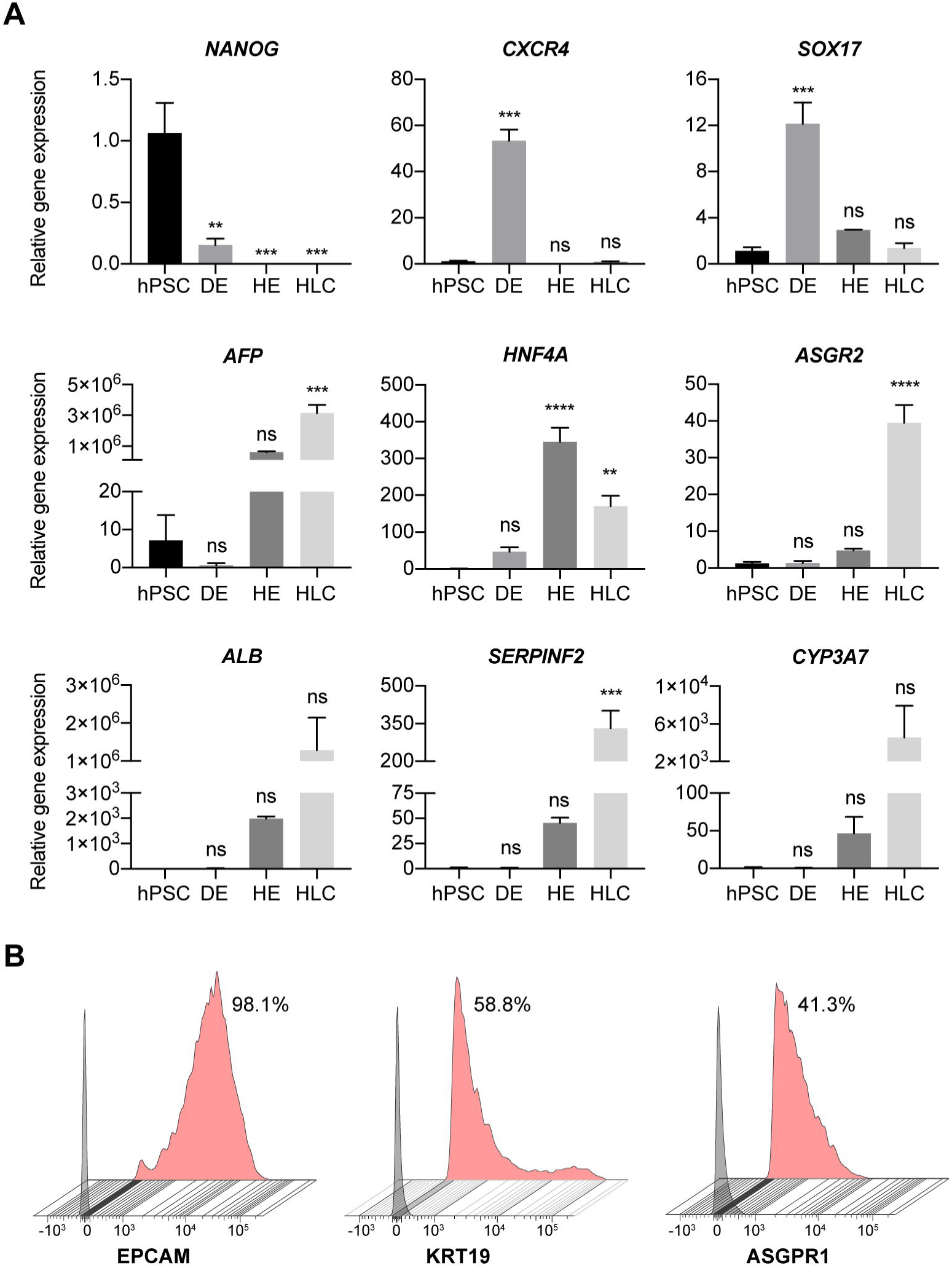
Phenotypical characterization of hepatic differentiated cells. **(A)** Differential expression of selected genes reveals progressive maturation of human pluripotent stem cells (hPSCs) to definitive endoderm (DE), hepatic endoderm (HE) and then hepatocytes (hPSC-Heps). Expression relative to the housekeeping gene, and normalized against the average expression of hPSCs; n=3 experiments, 1 cell line. Data are mean±SEM, ordinary one-way ANOVA followed by Dunnett post-hoc test to compare the mean of each group to hPSC expression. **p< 0.005; ***p< 0.0005; ****p< 0.0001; ns: nonsignificant. Data shown for cell line 1. **(B)** Flow cytometry analysis of surface marker expression on hPSC-Heps at day 21 of differentiation. Data shows the % of positive cells from the live cell population. Grey histogram represents FMO control used to establish the gate, red histogram represents stained hPSC-Heps. All flow cytometry analysis is representative of at least three independent experiments. Data shown for cell line 3.

Importantly the expression of *NANOG*, a key pluripotency gene, is significantly downregulated upon differentiation, and is obsolete at day 21, the stage in which hPSC-Heps have been produced. Furthermore, the expression of both *CXCR4* and *SOX17*, well established markers for both definitive endoderm and primitive endoderm, are found to peak and be significantly upregulated at day 7 of differentiation when compared to undifferentiated hPSCs; after this time-point the expression of both genes dissipates. To assess the correct differentiation into hPSC-Heps, we selected six major hepatic genes for investigation. By day 21 a significant increase in the relative expression of *AFP*, can be measured. *AFP* is the gene encoding alpha-fetoprotein, a major plasma protein produced be the developing liver and considered to be the fetal version of albumin [48]. Incidentally, expression of *ALB*, albeit not significant, is detected at days 14 and 21 of differentiation, with the expression being greatest at the later time-point. This provides evidence that at this stage the cell type produced is one that resembles a maturing hepatocyte. In addition, the relative gene expression for *HNF4A*, a hepatic transcription factor, as well as *ASGR2* (day 21 only), which encodes an asialoglycoprotein receptor isoform primarily found on liver cells, are significantly higher than that of undifferentiated or definitive endoderm cells. In the case of *HNF4A*, the relative gene expression is peaked at day 14, indicative that the hepatocyte-fate determination has occurred. Moreover, the expression for *SERPINF2*, the gene encoding the serpin alpha 2-antiplasmin which is secreted in plasma by hepatocytes, is significantly elevated after 21 days of differentiation. Finally, non-significant, but elevated expression of *CYP3A7* is detected in day 21 hPSC-Heps than the earlier time-points. This cytochrome P450 3A family isoform is predominately expressed in the developing liver, with the translated enzyme is involved in the metabolism of drugs, together with the synthesis of cholesterol, and various other lipids.

After validating the hepatic specific morphology and gene expression, we next sought to characterize the population profiles of the cells generated using our adapted protocol by performing flow cytometry analysis on day 21 hPSC-Heps (Figure 3B). The hepatic progenitor markers EpCAM (98.1%) and cytokeratin-19 (58.8%) were expressed on most cells analyzed. In addition to these hepatic progenitor markers, expression of the asialoglycoprotein receptor 1 (ASGPR1), an endocytotic cell surface receptor specific to adult hepatocytes, was detected on 41.3% of day 21 hPSC-Heps. The presence of hepatocyte progenitor markers and non-significant gene expression of *ALB* and *CYP3A7* displays the need for further maturation culture to produce a cell more closely resembling an adult hepatocyte.

### 2D maturation of hPSC-Heps

Having validated that cGMP-compliant hPSCs were able to generate cells resembling immature hepatocytes when cultured in our differentiation conditions, we next aimed to advance their hepatic maturation through culturing the cells within different culture model systems to challenge the clinical relevance of our protocols. We first seeded the day 21 hPSC-Heps onto collagen-1 coated tissue culture plastic, as it is an extracellular protein that has been shown capable of supporting the long-term culture, and liver-specific functions, of isolated adult hepatocytes [49]. Upon seeding, the hPSC-Heps recovered their polyhedral morphology within two days. It should be noted that if not seeded to confluency, then many cells do not remain viable in 2D culture on collagen-1, and those remaining do not proliferate, or go on to develop a mature phenotype.

After three weeks of maturation culture on collagen-1 coated tissue culture plastic, we performed immunofluorescent staining to assess the hepatic maturity and heterogeneity of the hPSC-Heps (Figure 4A). Firstly, none of the pluripotency or endodermal markers, such as OCT4 and CXCR4 are detected in the cultures, which is a good indicator that the conversion of stem cell to hepatic lineage cell was completed in our protocol. It is encouraging to observe hepatocyte-specific markers, for example the nuclear transcription factor, HNF4α and protease inhibitor, A1AT, to be abundantly expressed throughout the cultures. To further delineate the maturity of hPSC-Heps cultured on collagen-I coated 2D surfaces, we assessed the presence of hepatoblast specification (AFP, KRT19 and EpCAM) and hepatocyte specification (ALB and ASGPR1) markers respectively. We found that even though the expression of mature markers, such as ALB and ASGPR1, are prevalent across the culture, we still observed substantial regions of cells that expressed AFP, KRT19 and EpCAM. The perseverance of these progenitor markers reveals a potential limitation of 2D culture in terms of differentiating a fully adult-like hepatocyte. Of note, zona occluding 2 (ZO2), a component of tight junction proteins was highly expressed in 2D culture, highlighting the polyhedral morphology of the cells.

**Figure 4.**
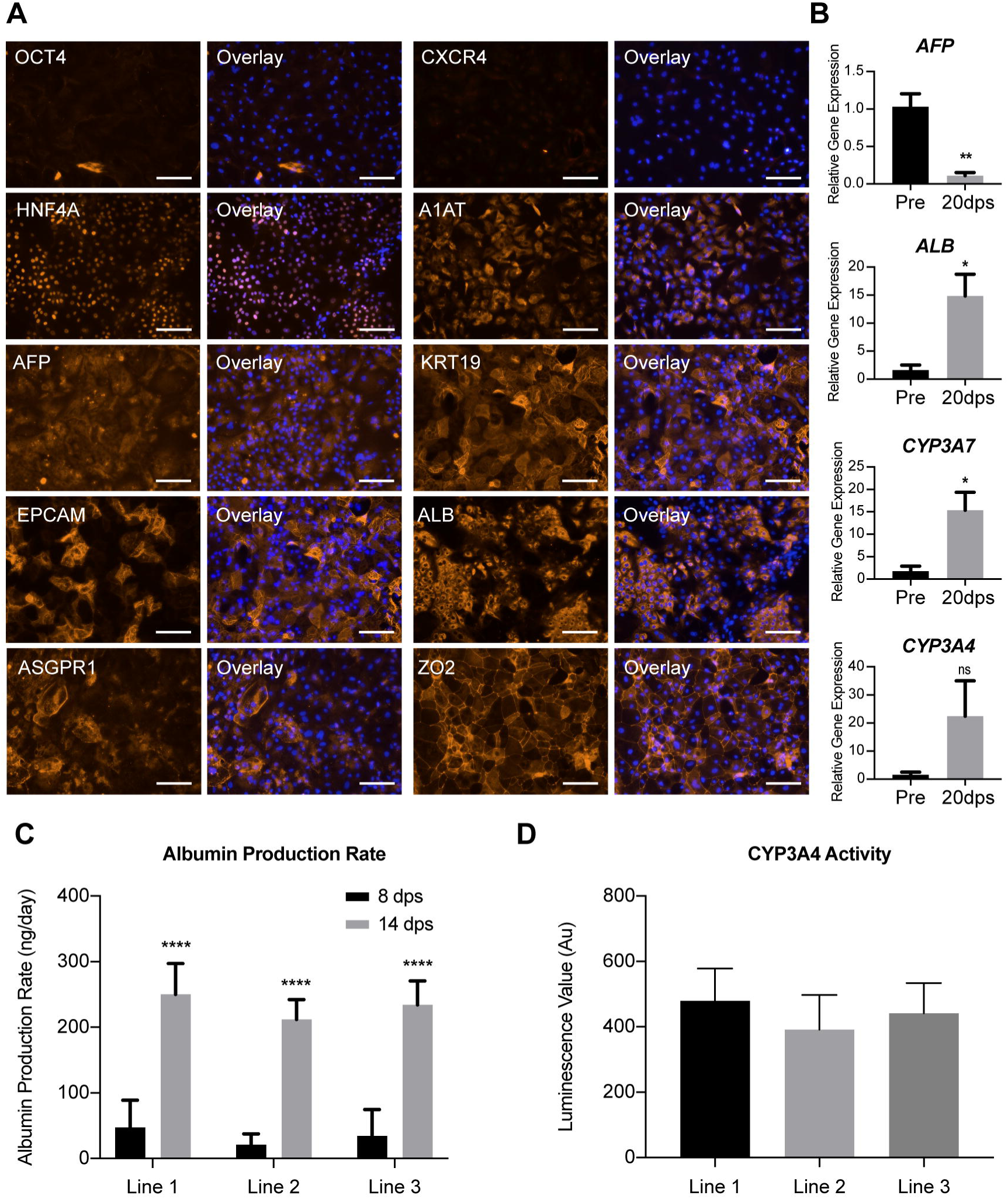
Characterization of hPSC-derived hepatocyte maturation in 2D model. **(A)** Immunofluorescent images revealing the transition of pluripotency (OCT4), endodermal (CXCR4) and hepatic specification (HNF4A, A1AT, AFP, KRT19 and EPCAM) and mature hepatic (ALB, ASGPR1 and ZO2) expression in hPSC-Heps after 2 weeks post-seeding on 2D model. Representative images selected from each of the 3 lines. Scale bar: 100μm. **(B)** Differential gene expression showing the relative expression of four key hepatic genes (*AFP, ALB, CYP3A7 and CYP3A4*) in pre-seeding (Pre) and 20 days post-seeding (20dps) into 2D model. Statistical significance determined by Student’s t-test (two-tailed); n=3 experiments, 1 cell line. Data are mean ± SEM, *p<0.05; **p< 0.005; ns: nonsignificant. **(C)** Albumin production rate of three cGMP-compliant cell lines 7 and 14 days post-seeding into 2D model; n=6 experiments per cell line. Data are mean ± SD, ****p<0.0001. **(D)** Cytochrome P450 3A4 enzyme activity of hPSC-Heps 3 weeks post-seeding into 2D maturation culture, n=3 (mean luminescence value [n=6] of 3 independent experiments).

For a broader evaluation of the maturation of hPSC-Heps in 2D, we compared the mRNA expression of cells 20 days post-seeding to that of the pre-seeding population (Figure 4B). A significant reduction in the level of *AFP* is achieved, whilst conversely, a significant enhancement in *ALB* expression occurs. A similar significant 15-fold increase is found in the relative expression of *CYP3A7* after the additional 20 days of culture. Interestingly, whilst enzyme activity was measurable in the previous assay, the gene expression of *CYP3A4* is elevated, but not significantly higher than that of the pre-seeding population. These mRNA expression results corroborate with our immunostaining observation that our cells are yet to achieve advanced maturity in this 2D model system.

To characterize the liver specific functions of the cells, we monitored the albumin production rate (Figure 4C and Supplementary data 3) and the metabolic activities of the cells over an extended period (Figure 4D). As a 2D monolayer, hPSC-Heps secreted albumin at levels detectable by ELISA after 8 days post-seeding. Once cultured for a further week, the amount of albumin protein measured was significantly higher and an indicator of a maturing phenotype. Moreover, the albumin production is comparable between the three lines assessed, with no significant differences measured after 8 or 14 days of maturation. Furthermore, when cultured for an additional week (21 days post-seeding), non-induced enzyme activity of cytochrome P450 3A4 can be detected (Figure 4D). Again, no significant difference between the different cGMP-compliant lines was measured, nor is activity detected in hPSC-Heps after one week of 2D maturation. The presence of CYP3A4 activity, which is initially absent from the liver of new-borns, and responsible for >50% of medicinal drug metabolism, is a clear indicator that the extended culture period results in matured hepatocytes [50,51].

### 3D maturation of hPSC-Heps within a PEG-DA-based scaffold suitable for biomedical applications

Having achieved considerable maturation on 2D collagen-1 coated tissue culture plastic, we proceeded to load day 21 hPSC-Heps into a more physiologically relevant model system for advanced hepatic maturation. We utilized a 3D macroporous PEG-DA hydrogel, known as an inverse colloidal crystal (ICC) scaffold, that aims to mimic the anatomy of native liver tissue. The ICC scaffold has uniform sized pores, interconnected in a hexagonal pattern, and we have previously demonstrated that human fetal liver-derived organoids can be generated through their culture within this model [52]. To facilitate the characterization of the morphogenic transformation of hPSC-Heps within this scaffold, we performed a series of immunofluorescence staining and constructed the 3D images using confocal microscopy. Firstly, we observed that upon seeding into the ICC hydrogel, hPSC-Heps establish cell-matrix interactions with the coated ECM protein and achieve confluency over the concave surface of the internal pores of the scaffold within 3-5 days post-seeding (Figure 5A). Then by one week, the cells self-assemble into mechanically stable interconnected clusters that resemble organoid structures for up to at least 3 weeks in culture. These distinct phases of morphogenesis were further illustrated with immunofluorescence imaging of beta-catenin (CTNNB1) and keratin 18 (KRT18) staining. Keratin 18 is the major intermediate filament protein in the liver, with a role in regulating glucose metabolism and modulating insulin signaling [53], while beta-catenin is the central constituent of canonical Wnt signaling and is implemented as a fundamental regulator in hepatic physiology and development [54].

**Figure 5.**
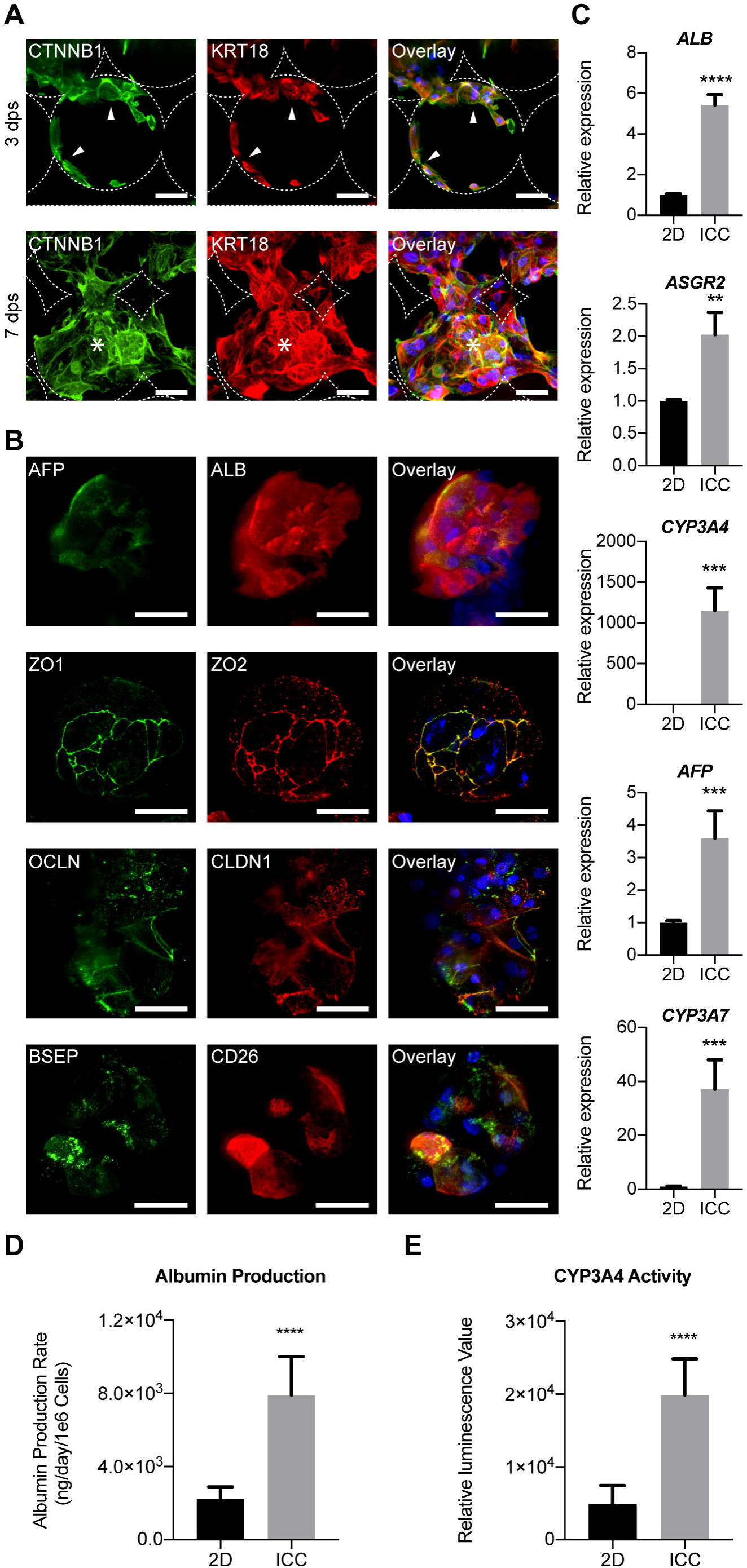
Characterization of hPSC-derived hepatocyte maturation within 3D ICC model. **(A)** Immunofluorescent confocal images of hPSC-Heps demonstrating two distinguished morphological phases inside the ICC scaffold. 3 days post-seeding an adhered lining across the hydrogel pores is observed, before the hPSC-Heps morph into interconnected 3D clusters from 7 days post-seeding onwards. Arrowheads indicate cells lining the ICC scaffold surface; asterisks represent cells forming 3D clusters. Scale bar, 100µm**. (B)** Immunofluorescent confocal images highlighting hepatic (AFP and ALB) and polarity (ZO1, ZO2, OCLN, CLDN1, BSEP and CD26) proteins known to be present in adult human hepatocytes. Scale bar, 100µm. Staining was performed on cell clusters after hPSC-Heps had been cultured for 2 weeks in 3D. **(C)** Real-time polymerase chain reaction showing the relative expression of 5 major hepatic genes (*ALB, ASGR2, CYP3A4, AFP* and *CYP3A7*); n=4 experiments, 1 cell line. **(D)** Albumin production rate of hPSC-Heps cultured in 2D vs ICC models; n=4 experiments, 1 cell line. **(E)** CYP3A4 basal activity of hPSC-Heps cultured in 2D vs ICC scaffolds; n=4 experiments, 1 cell line. Data are mean ± SD. Student’s t-test (two-tailed) analysis. **p<0.005; ***p<0.0005; ****p<0.0001. Data shown for cell line 1.

To further evaluate the phenotype of hPSC-Heps within this ICC culture model, we selected additional hepatic specific markers, along with markers of hepatocyte polarity, for broader evaluation by immunofluorescence imaging. Closer inspection on confocal micrograph reveals that the self-assembled interconnected clusters inside the ICC scaffold were consisting of a heterogeneous population, with AFP positive liver progenitor cells occupying the periphery of the cluster and surrounding the ALB positive mature hepatocytes at the core (Figure 5B). With most cells of the organoid-like clusters staining positive for cytoplasmic albumin, amid only few positive for AFP, this staining serves as confirmation that the seeded hPSC-Heps go on to develop a more mature phenotype. Moreover, correctly localized expression of the tight junction proteins ZO-1, ZO-2, occludin and claudin-1, bile-salt efflux pump (BSEP) and dipeptidyl peptidase-4 (CD26) are observed. These proteins are key components of bile canaliculi that form between the lateral faces of hepatocytes, and merge into bile ductules. The performed immunofluorescence staining provides validation of the liver-specific signature of the organoid-like structures generated.

We then performed a series of characterization to interrogate whether seeding hPSC-Heps into this 3D environment resulted in the acquisition of greater maturity compared to the 2D culture model. RT-PCR reveals that the expression of mature hepatic genes for proteins involved in biosynthesis (*ALB*), glycoprotein homeostasis (*ASGR2*) and metabolic functions (*CYP3A7* and *CYP3A4*) were significantly upregulated in the cells cultured within ICC scaffolds compared to those in 2D (Figure 5C). Notably, the expression of fetal hepatocyte associated AFP was significantly higher in the 3D culture, however this level is still significantly lower than in the day 21 cells initially seeded into the scaffold (Supplementary data 3).

Likewise, functional assays for albumin secretion (Figure 5D) and CYP3A4 enzyme activity (Figure 5E) showed significant improvements for hPSC-Heps matured within the ICC scaffold when compared to their 2D counterparts. The attainment and enhancement of these cellular functions are both major features of advanced hepatic differentiation, and their presence would be an unconditional necessity for any potential stem cell-derived therapy. Cumulatively, these data confirm that cGMP-compliant hPSC-derived hepatocytes can successfully be matured into a functional hepatic phenotype within a 3D, readily up-scalable, system. This combination of cGMP-compliant cells and a biocompatible scaffold not only provides a platform conducive to the further study of hPSC-Heps, but could also hold potential for future drug development and safety studies, and for assisting as a vehicle for cell transplantation.

### Generation of alginate encapsulated hepatocyte spheroids

To assess the broader translational potential of cGMP-compliant hPSC-derived hepatocytes for cell-based therapies aimed at ALF, we carried out microencapsulation of hPSC-Heps within alginate; a methodology that allows transplanted cells to be isolated from the recipient’s immune responses [55]. Successful encapsulation of hepatocytes within alginate hydrogels has been reported [56]. However, to reduce the possibility of cell death during the encapsulation process [57], we utilized an additional established culture model, spheroid culture [58]; known to both prolong viability [59] and phenotypes of hPSC-Heps [60] (Supplementary data 4).

To generate spheroids for encapsulation (Figure 6A), we utilized non-adherent microwell containing plates, called AggreWell^™^, which have previously been used to generate spheroids from hiPSCs [61]. We centrifuged single cell suspensions of hPSC-Heps into microwells to facilitate cell-cell interactions for spheroid formation in large readily scalable quantities (Figure 6B). The spheroids were left in culture for 7 days prior to encapsulation within 1.8% alginate microspheres. The 3D aggregates maintain their structure and uniformity as they are pumped through the microcapsule generator’s 0.22mm diameter nozzle at a rate of 10ml/hour. The cross-linking of alginate occurs in under 5 minutes within barium chloride (BaCL), with most microspheres containing a single spheroid. We then washed the collected microspheres in saline and subsequently placed back into HepatoZYME-SFM. To ensure that the viability of these hPSC-Heps was preserved throughout this process, we carried out live/dead immunofluorescence staining using fluorescein diacetate (FDA) and ethidium homodimer-1 (EthD-1) 6 hours after the encapsulation (Figure 6C). Confocal microscopy confirmed that both alginate- and non-capsuled spheroids contain minimal to no dead cells (EthD-1 stained nuclei), with close to all cells having hydrolysed FDA into fluorescent fluorescein. Furthermore, encapsulation does not impact the viability of hepatocyte spheroids placed back into further culture (Supplementary Data 5).

**Figure 6.**
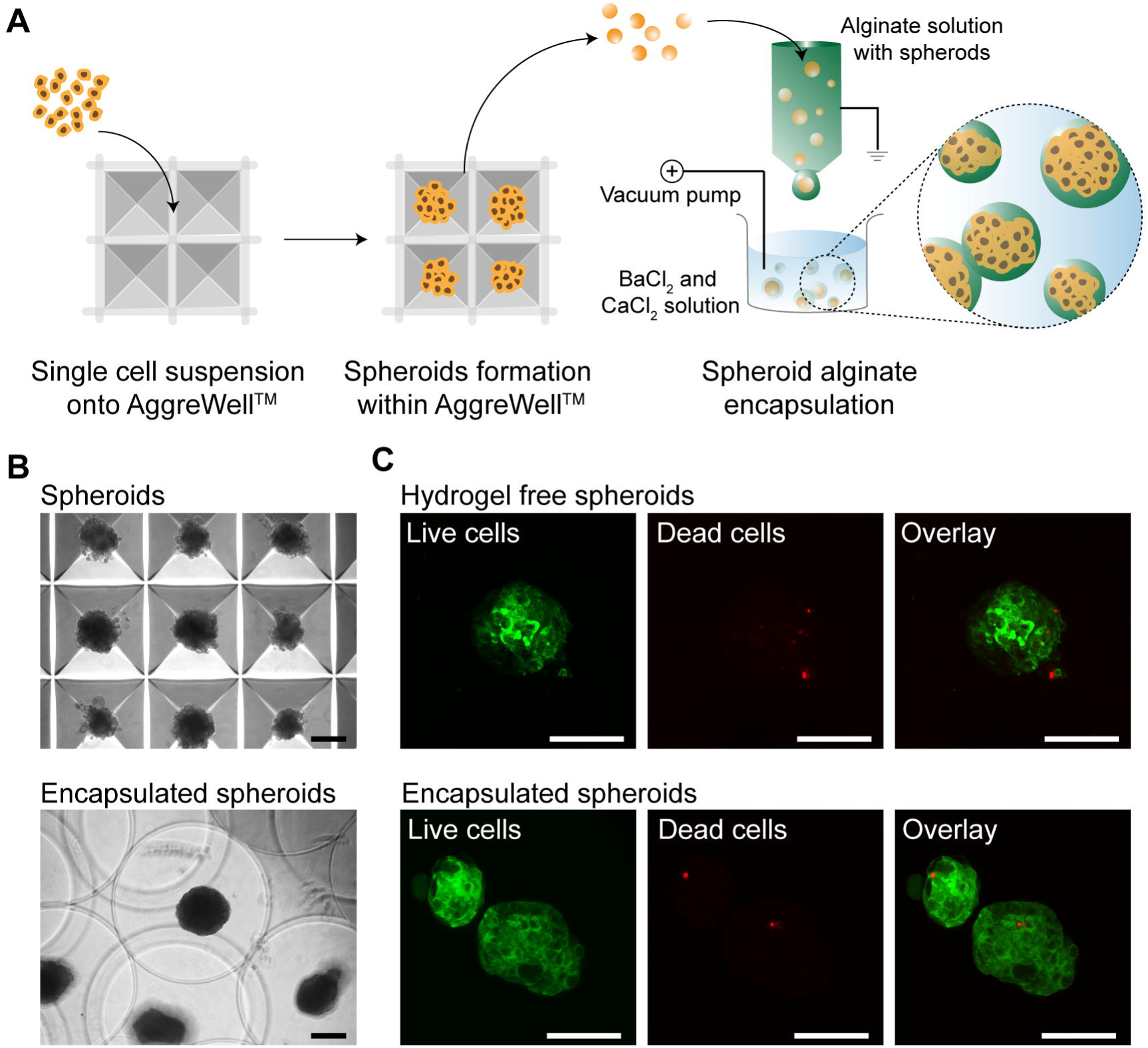
Generation of alginate encapsulated hepatocyte spheroids suitable for ALF bridging therapy. **(A)** Schematic illustrating the high throughout generation of uniform hepatocyte spheroids made up from around 250 hPSC-Heps using multi bioinert V-bottom microwells and electrostatic alginate microsphere encapsulation within a BaCl2 and CaCl2 solution bath. **(B)** Brightfield images showing hepatocyte spheroids inside AggreWell^™^ microwells and alginate microsphere encapsulated spheroids. **(C)** Confocal images revealing the live/dead staining of hPSC-Heps as hydrogel-free spheroids, and within alginate microspheres, 6-hours post encapsulation. Scale bars, 100µm.

In summary, we report that a library of cGMP-compliant hPSCs can be differentiated into hepatocytes by a chemically defined protocol, which is suitable for clinical implementation. We confirm that this development goes via an endoderm-like stage, before immature hepatocytes are obtained after three weeks. When matured by further culture in either 2D, or 3D models, these hPSC-Heps express proteins known to be present on adult hepatocytes, have improved hepatic gene expression, and go on to demonstrate key liver functions.

## Discussion

The manufacture of cGMP-compliant hPSC lines and optimization of translationally relevant differentiation technologies, is essential for clinical application of hPSCs [62]. We have, to our knowledge for the first time in this report, validated that a library of clinical grade cGMP-hPSC can successfully be differentiated into hepatocytes in a chemically defined protocol. Cells generated in this way demonstrated genes, proteins and hallmark functional characteristics of hepatocytes, but as shown by us and others previously [63], fell short of the benchmarks set by primary adult cells. By subsequently seeding these immature hepatocytes into bio-engineered 3D scaffolds fabricated from FDA approved material, we were able to drive the cells into liver tissue functionally approximated to the standard needed for clinical efficacy.

Following maturation within our 3D scaffold, elevated expression of key hepatic genes, such as *ALB*, *CYP3A7* and *CYP3A4* for example were all found to occur. CYP3A7 was first deemed as being exclusively expressed in the developing fetal liver [64], but it is now known to be present in up to 88% of adult livers [65,66]. This elevation in expression is of importance because cytochrome P450 enzymes are essential for the metabolism of numerous endogenous compounds and drugs. The CYP3A sub family specifically makes up 30% of the adult liver’s cytochrome P450 constituency [67] and metabolize half of marketed drugs [68]. Importantly, the level of CYP3A4 enzyme activity is also elevated in hPSC-Heps within 3D ICC culture when compared to 2D; as is the albumin production rate of the cells. It is also important to highlight that this scaffold can be readily scaled-up to hold greater numbers of cells – up to the billions required for human translation – and can be functionalized with different recombinant proteins, molecules or mechanical parameters. Cumulatively, these design features suggest this scaffold could be the ideal carrier for delivering cGMP-compliant hPSC-Heps into human patients.

Assuming delivery of suitable numbers of functionally optimized cGMP cells can be achieved as above, a further challenge for clinical application will be to deal with the potential allogenic immune rejection by the host post engraftment [69]. Alginate hydrogel microencapsulation provides hPSC-derivatives with a physical barrier from the recipient’s immune system, through enclosure within a naturally occurring anionic polymer. Alginate, typically obtained from brown seaweed, is considered ideal for biomedical applications due to its biocompatibility, low cost, and ease of gelation [70]. Numerous cell types have successfully been encapsulated within alginate, including mesenchymal stromal cells [71], pancreatic islets [72], and human hepatocytes [73], whilst optimized GMP grade alginate encapsulation protocols have already been established for the transplantation of human hepatocytes to provide metabolic function in patients with ALF [74]. By demonstrating cGMP-derived hepatic constructs remain viable post encapsulation within alginate hydrogel microspheres, we have therefore confirmed a potential road-map of hPSC-hepatocyte production appropriate for human use.

## Conclusion

We report here a library of clinical grade hPSCs manufactured under cGMP conditions amenable to reproducible hepatic differentiation, 3D culture and alginate encapsulation that may be suitable for human application.

## Acknowledgments

Generation of the GMP line LiPSC-GR1.1 was supported by the NIH Common Fund Regenerative Medicine Program, and reported in Stem Cell Reports. The NIH Common Fund and the National Center for Advancing Translational Sciences (NCATS) are joint stewards of the LiPSC-GR1.1 resource. We acknowledge Cell and Gene Therapy Catapult (London, UK) and Dr. Ricardo Baptista for the generation and provision of the CGT-RCiB-10 hiPSC line.

We thank the Nikon Imaging Centre at Kings College London for help with spinning disk confocal microscopy. We are grateful to the BRC Flow Cytometry Facility, KCH NHS Foundation Trust for advice and technical assistance.

## Disclosure of Potential Conflicts of Interest

The authors declare that there is no conflict of interest regarding the publication of this article

**Author Contributions**
Samuel J I Blackford: Concept and design, Collection and/or assembly of data, data analysis and interpretation, manuscript writing
Soon Seng Ng: Concept and design, Collection and/or assembly of data, data analysis and interpretation, manuscript writing
Joe M Segal: Collection and/or assembly of data
Aileen J F King: Collection and/or assembly of data
Jennifer Moore: Provision of study material or patients
Michael Sheldon: Provision of study material or patients
Dusko Ilic: Provision of study material or patients
Anil Dhawan: Concept and design, manuscript writing
Ragai Mitry: Concept and design, Collection and/or assembly of data, manuscript writing
S. Tamir Rashid: Concept and design, financial support, manuscript writing, Final approval of manuscript

## Acknowledgments of Grants

SJIB is supported by a GSTT BRC PhD award

STR is supported by an MRC Clinician Scientist Award (MGSBACR)

